# Direct detection of electrogenic polyamine transport

**DOI:** 10.64898/2026.10.01.755726

**Authors:** Sean R. Cleary, Cristina Moreno, Shubhra Srivastava, Pablo Miranda, Francesco Tadini-Buoninsegni, Miguel Holmgren

## Abstract

Polyamines are cationic metabolites that support critical cellular processes including transcription, translation, and cell proliferation. Although polyamine synthesis and transport are emerging therapeutic targets in human disease, the polyamine transport system remains poorly understood. ATP13A2-4 were recently identified as endolysosomal polyamine exporters, transporting putrescine (PUT), spermidine (SPD), and spermine (SPM) from the vesicular lumen to the cytosol. Despite interest in pharmacologically targeting these transporters, existing assays of transporter function rely on indirect or endpoint measures that cannot resolve transport as it occurs. Here, we report the first voltage-clamp recordings of polyamine transporter activity, directly detecting ATP13A2- and ATP13A3-mediated charge movement across cell membranes. When expressed in *Xenopus* oocytes, these transporters localized to the plasma membrane, enabling electrophysiological measurements. Polyamines induced inward currents dependent on transporter catalytic activity, and their dose-dependence correlated with ATPase assays. ATP13A2 currents were largest in response to SPM, whereas ATP13A3 currents were largest with PUT, consistent with their established substrate preferences. Polyamine-induced currents were blocked by the polyamine transport inhibitor AMXT-1501. These findings establish ATP13A2 and ATP13A3 as electrogenic polyamine pumps and demonstrate the utility of two-electrode voltage clamp as a label-free, transporter-resolved assay to complement other approaches in polyamine transporter-targeted drug discovery.

## Introduction

Polyamines are cationic metabolites that are required for a wide range of essential processes in all living cells. Specifically, polyamines play important roles in modulating transcription and translation, supporting autophagy, buffering mitochondrial reactive oxygen species, and regulating ion channel function (2-6). The eukaryotic polyamines putrescine (PUT), spermidine (SPD), and spermine (SPM) are produced as byproducts of amino acid metabolism. Their cytosolic availability is maintained through a combination of metabolic biosynthesis and intracellular transport (2). Although the metabolic pathways that synthesize polyamines have been well characterized and extensively investigated as therapeutic targets in multiple diseases (7), very little is known about the polyamine transport system that shuttles these metabolites from one cellular compartment to another (2).

A family of ATP-dependent transporters residing within endolysosomal membranes has recently been identified as major components of the polyamine transport system in mammalian cells (8-11). ATP13A2, ATP13A3, and ATP13A4 are members of the P5B-ATPase family that mediate ATP-dependent export of polyamines from the endolysosomal lumen to the cytosol (8, 9, 11). Extracellular polyamines bind cell surface proteoglycans and enter the endosomal system by endocytosis, where they encounter these transporters (12). The three polyamine-transporting ATPases exhibit differential tissue specificity. While ATP13A2 and ATP13A3 are thought to be ubiquitously expressed, ATP13A2 expression is elevated in the central nervous system (CNS). The expression of ATP13A4 is more restricted to specific tissues and cell types, including endocrine glands and neuronal astrocytes. An additional P5B transporter ATP13A5, expressed in CNS pericytes in the brain, spinal cord, and retina, is also predicted to transport polyamines (13, 14). These transporters also exhibit distinct yet overlapping subcellular distribution along the endosomal pathway. While ATP13A2 mainly resides in late endosomes and lysosomes, ATP13A3 is in early endosomes and recycling vesicles, and ATP13A4 exhibits a broader distribution throughout the endosomal system (15).

Polyamine metabolism and transport have emerged as potential therapeutic targets in diverse human disease states (16-20). Because intracellular polyamines support cell proliferation, strategies that disrupt polyamine homeostasis have been extensively explored in cancer. Difluoromethylornithine (DFMO), an inhibitor of polyamine biosynthesis, was FDA-approved to reduce the risk of relapse in high-risk neuroblastoma, and its combination with the polyamine transport inhibitor AMXT-1501 is being evaluated in clinical trials as a strategy to deplete intracellular polyamines in multiple forms of cancer (18-22). Dysregulated polyamine homeostasis has also been implicated in neurodegenerative disease (8, 23, 24). Decreased levels of SAT1, the rate-limiting enzyme in polyamine catabolism, were found to be a hallmark of idiopathic Parkinson’s disease, linking altered polyamine metabolism to disease pathogenesis (24). Moreover, loss-of-function mutations in *ATP13A2* are the cause of Parkinson’s disease type 9 (PARK9), a monogenic form of Parkinson’s, directly linking reduced intracellular polyamine transport to neurodegeneration (2, 3, 8).

Despite the interest in targeting polyamine transporters in disease, the human polyamine transport system is still being elucidated (2, 13, 25, 26). As a result, critical tools to study polyamine transporters, like direct activity assays and selective small molecules, remain underdeveloped. Presently, two principal approaches are used to characterize the function of ATP-dependent polyamine transporters: (I.) measurement of polyamine-stimulated ATP hydrolysis and (II.) determining whole cell uptake of labeled-polyamines (27). However, these approaches can only provide indirect or endpoint readouts that do not resolve individual transporter activity in real time. In contrast, electrophysiology records transporter-mediated charge movement as it occurs, providing a continuous, label-free readout of electrogenic transport. This could provide access to dynamic features of the transport process and enable mechanistic dissection of how biochemical, pharmacological, and pathological perturbations alter transporter function. Electrophysiological techniques have been used for decades to study other electrogenic P-type ion pumps, like the Na^+^/K^+^ ATPase (28). Yet, the electrophysiological analysis of endolysosomal transporters like ATP13A2 is challenging, as intracellular organelles are generally inaccessible by standard approaches. Here, we report the functional expression and detection of polyamine transporter activity through the direct measurement of transporter-mediated charge movement across the plasma membrane of oocytes from *Xenopus laevis*. These transporter-resolved polyamine currents introduce a powerful platform to mechanistically study polyamine transporters, screen targeted therapies, and evaluate disease variants in future studies.

## Results

### Expression of ATP13A2 in the cell surface of Xenopus oocytes

ATP13A2 localizes to lysosomes and endosomes in mammalian cells, making it inaccessible to whole cell electrophysiological recordings. To subvert this barrier, past researchers have leveraged *Xenopus laevis* oocytes which exhibit promiscuous trafficking of membrane proteins to the plasma membrane. Indeed, past studies have demonstrated that other endolysosomal transporters can localize to the cell surface in *Xenopus* oocytes (29-31). Similarly, expressing a fluorescent tagged ATP13A2 (ATP13A2-GFP) in oocytes revealed a population of ATP13A2 with apparent plasma membrane localization (**Fig. 1A, Fig. S1**). To confirm plasma membrane localization of ATP13A2, we expressed an ATP13A2 construct with a hemagglutinin (HA) interdomain tag inserted after Val1030 in a long unstructured loop between the transmembrane helices TM7 and TM8 (ATP13A2-HA) (1). This loop, which normally faces the endolysosomal lumen, is reoriented outward during vesicular membrane fusion (**Fig. 1B**), exposing it to the extracellular space. Chemiluminescent immunostaining with an anti-HA antibody revealed that oocytes injected with ATP13A2-HA cRNA showed significantly increased luminescent labeling compared to oocytes injected with unlabeled ATP13A2 cRNA (*p* = 6.7E=16) or water (*p* = 9.9E-16) (**Fig. 1C,D**). This indicates that the HA-tag is extracellularly exposed, supporting that ATP13A2 is indeed present at the plasma membrane of oocytes.

**Figure 1.**
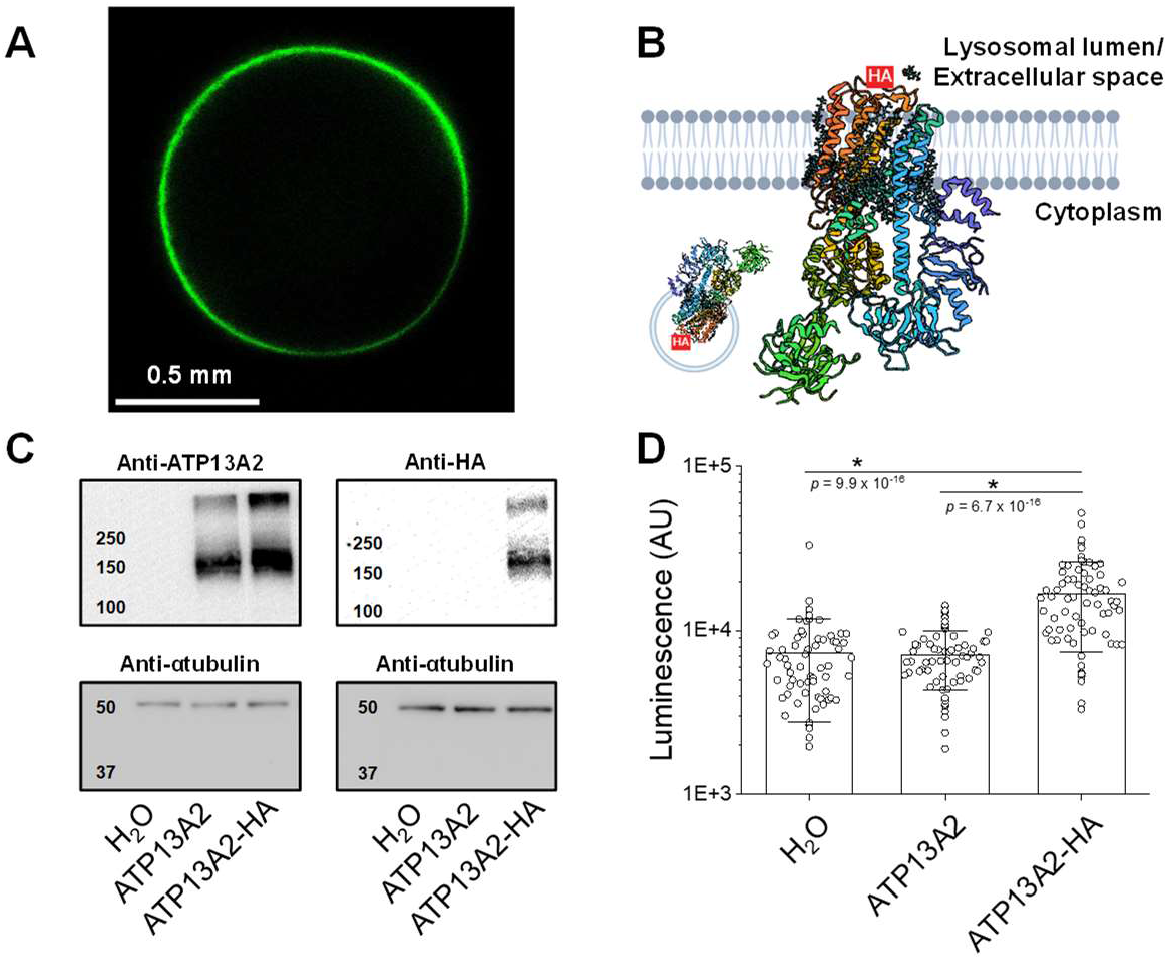
A population of ATP13A2 mislocalizes to the plasma membrane of *Xenopus* oocytes. **A)** Confocal image of a population of ATP13A2-GFP localized to the plasma membrane of *Xenopus* oocytes. **B)** A cartoon schematic diagram of ATP13A2-HA membrane orientation (PDB: 7M5X) (1). **C)** Western blot detects expression of ATP13A2 and ATP13A2-HA. **D)** Chemi-luminescent staining of an extracellular facing HA-tag inserted in ATP13A2 (H_2_O *n* = 67, ATP13A2 *n* = 66, ATP13A2-HA *n* = 72). Differences determined by 1-way ANOVA with Dunn Sidak *post-hoc* (*F* = 54.1, *p* = 1.6 x 10-19).

### Functional measurements of ATP13A2 ionic currents

Plasma membrane localization of the ATP13A2 in *Xenopus* oocytes enables the application of classical electrophysiology methods to study its function, provided it produces detectable electrogenic currents. At present, it remains uncertain whether ATP13A2 transport is expected to be electrogenic or how changes in membrane voltage may affect its transport cycle. We used the two-electrode voltage clamp (TEVC) approach to probe oocytes injected with ATP13A2 cRNA for polyamine-induced currents. While the oocytes were continuously perfused with extracellular bath solutions (± 1 mM SPM), a 10 mV voltage-step protocol from -190 to +60 mV (**Fig. 2A**) was applied every 1 minute (P1-P7, **Fig. 2B**), with -190 mV representing the lower voltage limit of the amplifier and the holding potential maintained at 0 mV. This experimental design permits assessment of the stability of the preparation over time by subtracting the current at each voltage between protocols performed with similar solution compositions (e.g. P2-P1, *Time Control*, **Fig. 2C**).

**Figure 2.**
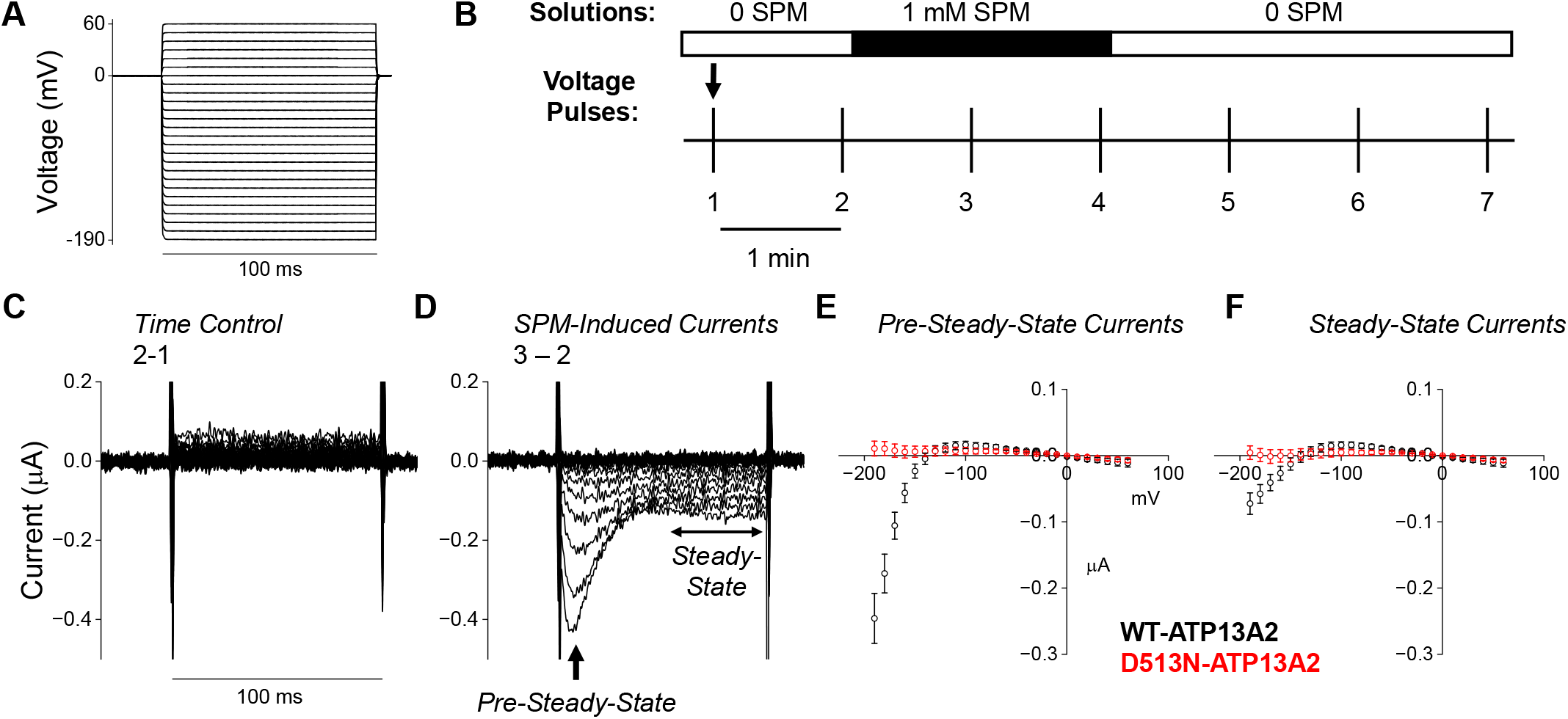
Functional measurements of polyamine-induced ATP13A2 activity-dependent charge movement. **A**,**B)** A voltage protocol of 100 ms pulses from -190 to +40 mV (10 mV step, holding potential = 0 mV) was applied to oocytes every 1 min while perfusing with bath solution ± 1 mM SPM. **C)** No inward (negative) current detected in ATP13A2 expressing oocytes in the absence of SPM. **D)** Subtracting steady-state currents detected in the absence of SPM from currents elicited in its presence revealed a SPM-induced current consisting of a pre-steady-state current followed by a smaller steady-state current. **E**,**F)** Voltage dependence of the pre-steady-state (E) and steady-state currents (F) as represented in D (WT *n* = 14, D513N *n* = 7).

Similarly, SPM-induced currents were obtained by subtracting currents in the absence of SPM from those obtained in its presence (e.g. P3-P2, **Fig. 2D**). Interestingly, as voltages became more negative, SPM-induced inward currents were detected exhibiting a pre-steady-state component followed by smaller steady-state currents, both with similar voltage dependence (**Figs. 2E** and **2F**). The experimental design also allows us to assess whether these polyamine-induced currents reverse as the polyamine is washed away (e.g. P4-P7). Indeed, the steady-state currents were completely reversed within 3 minutes of perfusion with bath solution (**Fig. S2**). These currents were not detected in uninjected oocytes or oocytes expressing a catalytically incompetent mutant ATP13A2 (D513N) (**Figs. 2E, 2F**, and **Fig. S2**). This mutation neutralizes a conserved aspartate required for P-type ATPase autophosphorylation, abolishing the current without altering expression (**Fig. S2D**), indicating that these currents depend on ATP13A2 enzymatic activity.

Next, we characterized the SPM dose-dependence of the ATP13A2 currents induced over a range of SPM concentrations during voltage steps from 0 to -190 mV (**Fig. 3A,B**). Increasing SPM concentration resulted in progressively larger currents, with responses detectable at concentrations as low as 10^-7^ M and saturating in the millimolar range (**Fig. 3C,D**). Interestingly, the maximal amplitude of the pre-steady-state currents appears to have a biphasic response to increasing SPM concentrations (**Fig. 3C, Fig. S3)**. A Hill fit of the dose-dependences of these two currents revealed that the ascending phase of the pre-steady-state currents had a significantly lower apparent *K*_*m*_ (4.5 ± 2.6 µM) with respect to the steady-state current response (43.3 ± 11.2 µM, *p* = 0.004, paired two-tailed t-test) (**Fig. S3**), indicating that these currents may correspond to distinct processes within the polyamine transport cycle.

**Figure 3.**
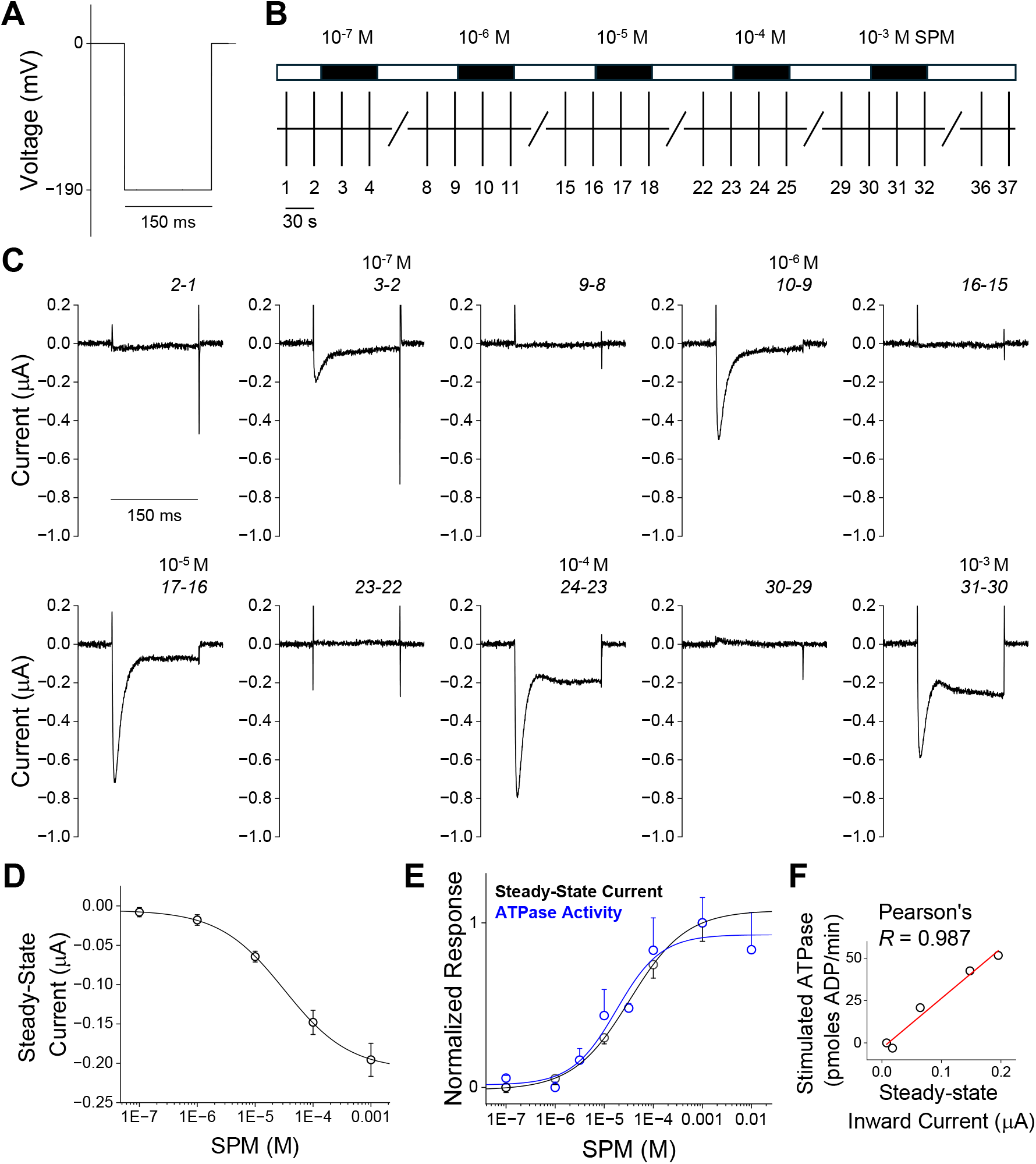
Characterization of dose-dependence of SPM-induced ATP13A2 currents. **A, B)** Voltage pulses from 0 to -190 mV for 150 ms (holding potential = 0 mV) were applied to oocytes every 30 s while perfusing with bath solution ± SPM. **C)** Representative current traces showing concentration-dependence of SPM-induced ATP13A2 currents, with complete washout between doses. Traces show subtracted currents, with labels indicating the corresponding pulse numbers in *B*. **D)** Dose-dependent activation of steady-state currents in response to increasing SPM concentration (*n* = 9). **E)** The dose dependence of SPM-induced steady-state currents is comparable to SPM-stimulated ATPase activity (*n* = 3). **F)** SPM-induced steady-state current correlates strongly with SPM-stimulated ATPase activity over a range of SPM concentrations (as in E) (Pearson’s *R* = 0.987).

To compare these electrophysiological responses with established biochemical measures of ATP13A2 activity, we measured the SPM-dependent activation of ATP13A2 ATPase activity in lysosomes isolated from HEK293T cells overexpressing ATP13A2-GFP (**Fig. S4**). The ATP13A2-GFP construct exhibits SPM-induced currents, indicating the function of the tagged construct is preserved (**Fig. S5**). A Hill fit of ATP13A2 ATPase activity gave an apparent *K*_*m*_ similar to that of the steady-state current (33.4 ± 12.7 µM). Moreover, ATP13A2 ATPase assays and steady-state currents gave very similar dose-response relationships (**Fig. 3E**) that correlated strongly with one other (Pearson’s *R* = 0.987) (**Fig. 3F**). These results further establish that the steady-state currents likely reflect polyamine transport and are tightly coupled to ATP hydrolysis within the ATP13A2 enzymatic cycle. Since the pre-steady-state currents respond with substantially higher affinity than the steady-state currents, these transient signals may arise from early events that preceded polyamine transport.

### Polyamine substrate-dependence of ATP13A2 and ATP13A3 currents

To assess whether polyamine-induced currents reflect the known substrate preference of ATP13A2 for SPM over other polyamines, we compared currents induced by the polyamines PUT (2+), SPD (3+), and SPM (4+) during voltage steps from 0 to -190 mV (**Fig. 4A-C**). Indeed, consecutive measurements within the same oocyte revealed that the steady-state currents increased from PUT to SPD to SPM (**Fig. 4D**). Notably, each unit increase in polyamine charge (+1) produced a linear increase in the average ionic current detected (**Fig. 4E**). This correlation suggests that the steady-state currents reflect the known selectivity of ATP13A2 for SPM. Although all polyamines also elicited pre-steady-state currents, the amplitudes of these pre-steady-state signals varied greatly and did not scale with the polyamine charge (**Fig. S6**). Moreover, subsequent experiments measured currents induced by these polyamines over a range of voltage steps from -190 to +60 mV, revealing similar voltage dependences for all three amines (**Fig. S6**).

**Figure 4.**
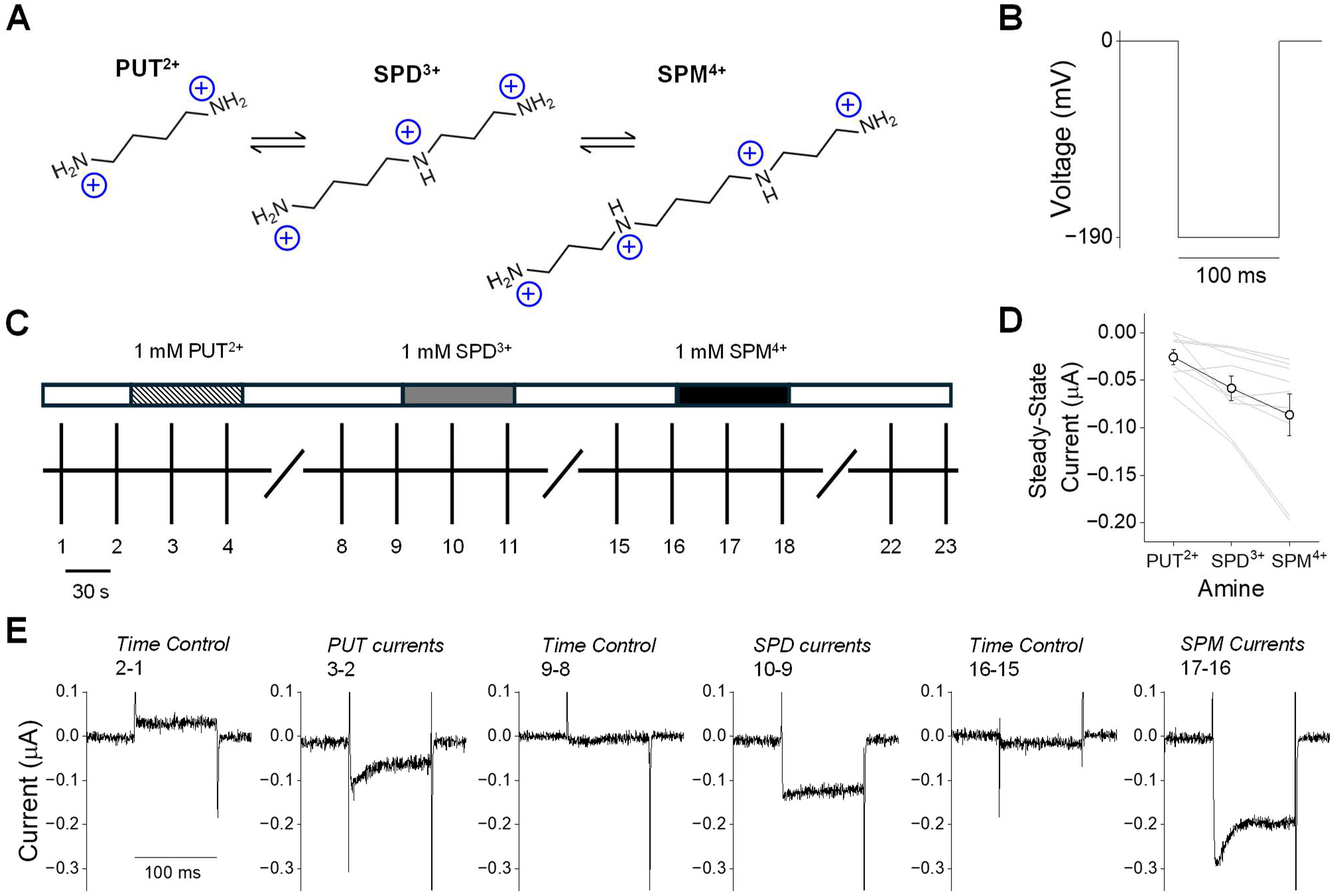
Characterization of substrate-dependence of ATP13A2 currents. **A)** Chemical structures of polyamines putrescine (PUT), spermidine (SPD), spermine (SPM). **B**,**C)** Voltage pulses of -190mV for 100 ms (holding potential = 0 mV) were applied to oocytes every 30 s while perfusing with bath solution ± 1mM SPM, SPD, or PUT. **D)** Larger currents were elicited by 1 mM SPM (+4) than SPD (+3) or PUT (+2) (*n* = 9). **E)** Representative current traces showing substrate-dependent activation of ATP13A2 currents, with complete washout between varying polyamines. Traces show subtracted currents, with labels indicating the corresponding pulse numbers in *C*.

ATP13A3 is a related endosomal polyamine transporter reported to preferentially export PUT over SPM to the cytoplasm (9). To assess its potential for electrogenic polyamine transport, we injected *Xenopus* oocytes with human ATP13A3 cRNA and tested for currents induced by 1 mM PUT, SPD, or SPM. Since ATP13A2 and ATP13A3 may have differing voltage dependences, we probed these cells for currents over a range of voltage potentials from -190 mV to +60 mV (**Fig. 5A,B**). Once more, increasingly negative potentials revealed novel polyamine-induced currents in ATP13A3 injected oocytes (**Fig. 5C-G**). In contrast to ATP13A2, the largest current amplitudes were induced by PUT and decreased when switching to SPD and to SPM (**Fig. 5C**). Although their amplitudes differed, each amine activated current within a similar voltage range (**Fig. 5D-F, Fig. S7**). No current was detected from oocytes expressing a catalytically incompetent mutant ATP13A3 (D498N), indicating that these currents depend on ATP13A3 catalytic activity. The preference of ATP13A3 currents for PUT is consistent with its established substrate selectivity, further supporting our current measurements as a direct readout of polyamine transport.

**Figure 5.**
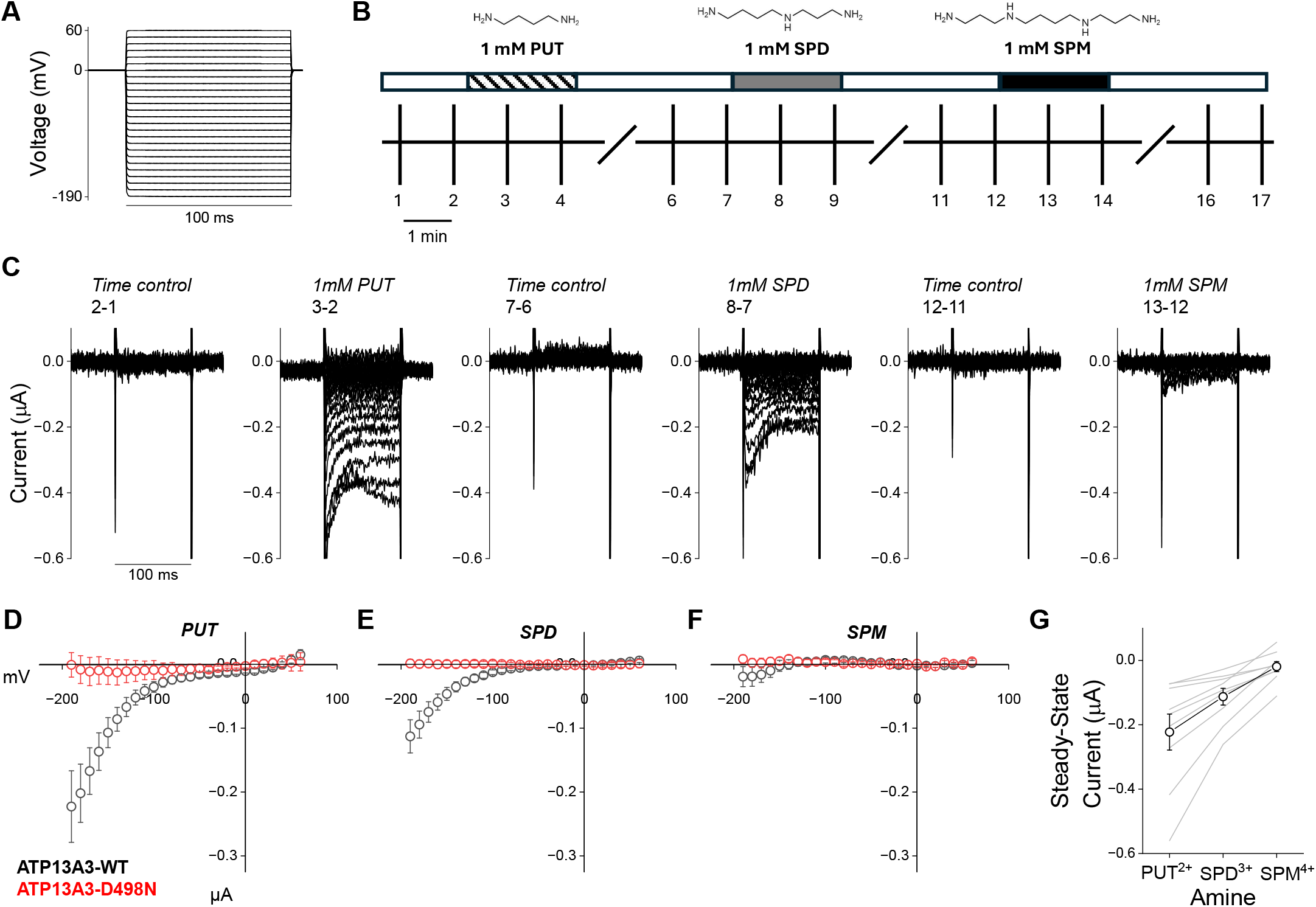
Characterization of substrate-dependence of ATP13A3 currents. **A**,**B)** A voltage protocol of 100 ms pulses from -190 to +40 mV (10 mV step, holding potential = 0 mV) was applied to oocytes every 1 min while perfusing with bath solution ± 1 mM PUT, SPD, and SPM. **C)** Representative current traces showing substrate-dependent activation of ATP13A3 currents, with complete washout between varying polyamines. Traces show subtracted currents, with labels indicating the corresponding pulse numbers in *B*. **D-F)** Voltage dependences of steady-state currents elicited from ATP13A3-WT injected oocytes (*black*) by 1mM PUT (*D*), SPD (*E*), and SPM (*F*) as shown in *C* (*n* = 9) compared to currents from oocytes injected with ATP13A3-D498N (*red*). **G)** Larger steady-state currents were elicited at -190 mV by 1 mM PUT than SPD or SPM (*n* = 9).

### Pharmacological inhibition of polyamine transport currents by AMXT-1501

AMXT-1501 is a small molecule competitive inhibitor of polyamine transport (32) and was recently shown to block activity of polyamine transporting P5B-ATPases, ATP13A2 and ATP13A3 (33, 34). The AMXT-1501 molecule chemically consists of a spermine molecule conjugated to a long lipophilic group (**Fig. 6A**) (32). To assess the effect of AMXT-1501 on our ATP13A2 polyamine currents, we monitored the SPM-induced currents in response to a series of 100 ms voltage steps from 0 to -190 mV (**Fig. 6B,C**). We began the experiments by establishing the 100 μM SPM-induced current (P3-P2, **Fig. 6D**). Then, we perfused a bath solution containing 100 μM SPM and 100 nM AMXT-1501 to obtain the AMXT-1501 sensitive current (P4-P8, **Fig. 6D**). The AMXT-1501 sensitive component of the currents reflected a similar amplitude to currents induced by SPM, indicating that AMXT-1501 is a potent inhibitor of ATP13A2 transport (**Fig. 6E**). To test the reversibility of AMXT-1501 inhibition of SPM currents, we perfused the oocytes with bath solution for 5 minutes following AMXT addition and then reintroduced 100 μM SPM. SPM-induced currents were not recovered after this period of perfusion (P19-P18, **Fig. 6D,E**), suggesting that AMXT-1501 inhibition is irreversible within this time frame. We also found that AMXT-1501 effectively blocked ATP13A3 currents induced by 100 µM PUT (**Fig. S8**). Interestingly, a higher concentration of AMXT-1501 (1 µM) was required to completely block the ATP13A3 current at this concentration of PUT, suggesting this inhibitor may be less selective for ATP13A3 compared to ATP13A2 (**Fig. S8**).

**Figure 6.**
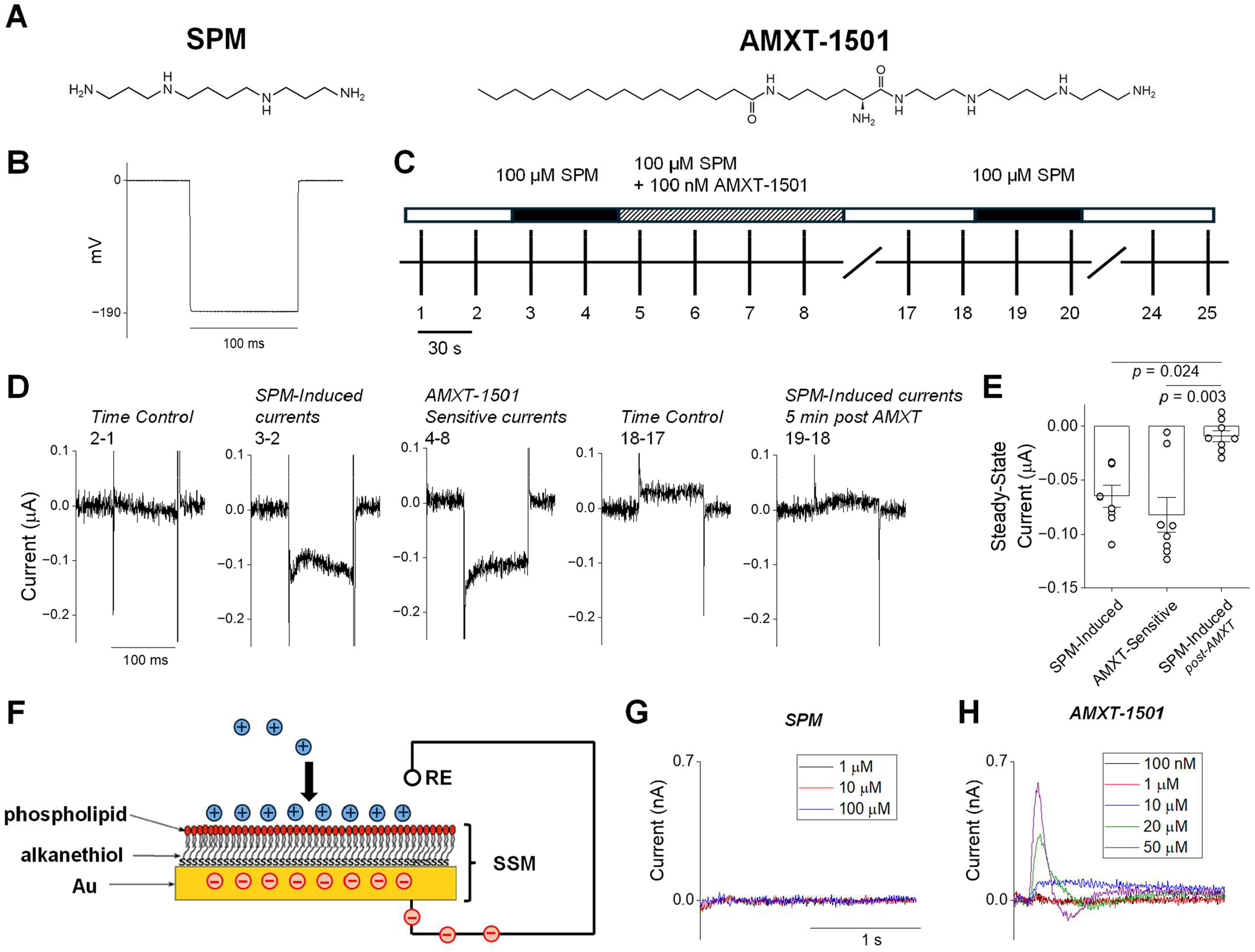
Competitive inhibition of ATP13A2 polyamine transport by AMXT-1501. **A)** Chemical structures of SPM and AMXT-1501. **B**,**C)** -190 mV, 100 ms depolarizations (holding potential = 0 mV) were applied to oocytes every 30 s while perfusing with bath solution or solution with 100 µM SPM ± 100 nM AMXT-1501. **D**,**E)** AMXT-1501 completely abolished SPM-induced currents. SPM-induced currents were not recovered after removing AMXT-1501 and perfusing with bath solution for 5 min (*n* = 8). Lines and error bars represent mean ± SEM and data were analyzed by Kruskal-Wallis with Dunn’s *post*-*hoc* test (χ^2^ = 12.2, *p* = 0.002). **F)** Schematic diagram of the interaction of cationic molecules (*blue spheres*) with the solid supported membrane (SSM). The SSM behaves like an electrical capacitor within the electrical circuit. If cations bind to the SSM membrane, electrons (*red spheres*) must flow through the external circuit towards the gold surface to compensate for the build-up of positive charge on the membrane. A capacitive current signal is therefore generated which is measured by the device. RE is the reference electrode. **G, H)** Current traces were recorded after SPM (*G*) and AMXT-1501 (*H*) concentration jumps on two different SSM membranes. The concentrations of SPM and AMXT-1501 are indicated in the panels. Each trace was obtained by averaging five consecutive measurements.

A plausible explanation for this low reversibility is that the lipophilic group of AMXT-1501 interacts with lipid membranes (**Fig. 6A**) (32). To test this possibility, we used solid supported membrane (SSM) electrophysiology. The SSM, consisting of an alkanethiol/phospholipid bilayer supported by a gold electrode (35), represents a convenient experimental model of a biological membrane. Such a bilayer is characterized by high mechanical stability so that solutions can be rapidly exchanged at the surface (36). SSM-based electrophysiology was recently employed to examine the membrane binding of the hydrophobic ions tetraphenylborate and tetraphenylarsonium, demonstrating the utility of this technique for detecting membrane interactions of charged molecules (37). The detection of ion binding to the SSM membrane relies on the principle that the SSM behaves like an electrical capacitor within the electrical circuit. When membrane-binding ions are introduced into the system above the SSM surface, this is equivalent to switching on a DC power supply, providing ions to charge up the membrane capacitor. This causes a transient capacitive current to be induced in the circuit (**Fig. 6F**, see *Methods*). The SSM method is clearly applicable to studies of the membrane binding of any charged compound, such as the positively charged molecules SPM and AMXT-1501. We therefore performed control experiments to investigate potential binding of SPM to the SSM membrane. SPM concentration jumps (from 1 to 100 µM) on the SSM did not generate a detectable current (**Fig. 6G**), suggesting that SPM does not interact with the membrane. We then carried out concentration jumps of AMXT-1501 from 100 nM to 50 µM on the SSM. As shown in **Fig. 6H**, a positive current was detected for AMXT-1501 ≥ 5 µM, indicating the binding of positive charge to the SSM. The amplitude of the current signal increased as the concentration of AMXT-1501 increased (**Fig. 6H** and **Fig. S9**). We therefore concluded that AMXT-1501 can interact with the supported membrane. In particular, binding of AMXT-1501 to the SSM membrane likely occurs via the lipophilic group of the AMXT-1501 molecule (**Fig. 6A**). It is worth noting that a current component of rather small negative amplitude could be observed at a high AMXT-1501 concentration (50 µM, **Fig. 6H**). We found that the negative current component decreased in amplitude when the AMXT-1501 jumps were performed from higher to lower concentration (**Fig. S9**). Such a negative current response might be attributed to a small number of positive charges moving away from the SSM membrane. We speculate that jumps of AMXT-1501 at increasing concentration might cause a progressive accumulation of positive charges on the SSM membrane, which eventually results in partial unbinding of some AMXT-1501 molecules due to electrostatic repulsion. Taken together, the results reveal new insights into the mechanism, selectivity, and reversibility of this small molecule inhibitor of polyamine transport.

## Discussion

Polyamines are critical cellular metabolites that are tightly regulated through a combination of synthesis, degradation, and transport (2). Polyamine transporters have emerged as potential therapeutic targets in a range of disease states, including infectious disease and cancer (16-20); yet, the activity of these transporters, particularly ATP13A2-4, has primarily been evaluated using biochemical or endpoint transport assays that do not resolve transporter activity in real time. Here, we establish voltage-clamp electrophysiological recordings as a direct approach to measure transport-associated currents from ATP13A2 and ATP13A3 in real time. To our knowledge, these represent the first electrophysiological recordings of active polyamine transport and demonstrate that both of these transporters are electrogenic. These measurements represent a powerful platform to study mechanisms of polyamine transporter function in live cell membranes.

ATP13A2 and ATP13A3 are members of the P-type ATPase family of transporters such as the Na^+^/K^+^-ATPase. Unlike ion channels, which allow ions to permeate down their electrochemical gradients at rates near diffusion, these active transporters move only a few ions per second at rates limited by their enzymatic cycles, making their electrogenic currents difficult to detect experimentally. This presents a particular challenge for eliciting measurable currents from endolysosomal transporters because the transporter population is divided across numerous small, electrically independent compartments. In our approach, detection of polyamine transporter currents was enabled through plasma membrane localization of these transporters in *Xenopus* oocytes. The exceptionally large membrane area permits expression of a large population of transporters that can be simultaneously subjected to defined membrane potentials, enabling their collective activity to be detected as measurable macroscopic currents (**Fig. 1**). Importantly, plasma membrane localization preserves the membrane orientation of ATP13A2 and ATP13A3 such that the extracellular milieu corresponds to the luminal side of endocytic membranes. Therefore, the polyamine-induced currents reflect the export of polyamines from endocytic compartments to the cytosol, consistent with the physiological function of these transporters (2).

These polyamine-induced currents establish that ATP13A2 and ATP13A3 mediate electrogenic polyamine transport. Because polyamines carry multiple cationic charges per molecule and induce an inward net charge movement, we can confidently conclude that the transported polyamines contribute substantially to this net charge displacement. Consistent with this interpretation, the currents exhibit the established substrate preferences of ATP13A2 for SPM (**Fig. 4**) and ATP13A3 for PUT (**Fig. 5**). However, it remains entirely possible that other ions could also contribute to the measured currents as co- or counter-transported substrates. Several other ions have been proposed as potential substrates of ATP13A2 including protons, potassium, calcium, manganese, and zinc (1, 38-40). Future studies could evaluate the impact of ionic substitutions on these currents. One important limitation of the two-electrode voltage clamp oocyte approach is substrate control is limited to the extracellular compartment. More advanced electrophysiological techniques like the cut-open vaseline gap approach could enable control over the intracellular compartment (41), enabling a more complete biochemical evaluation of how extracellular and intracellular ions affect the activity and net charge movement of these transporters.

In addition to providing a direct, real-time readout of polyamine transport, these measurements also revealed pre-steady-state currents coupled to the catalytic activity of the enzymes. The pre-steady-state currents decay during the pulse into the more sustained steady-state currents (**Fig. 2**) and exhibit notably higher apparent SPM affinity than the steady-state currents of ATP13A2 (**Fig. 3C** and **Fig. S3**). Together these distinct kinetics and concentration dependence are consistent with the pre-steady-state currents reflecting partial reactions of the enzyme associated with substrate binding and/or early conformational transitions. Similar pre-steady-state currents have been observed in other transporters including the Na^+^/K^+^-ATPase (42, 43). Still, more work is needed to evaluate which partial reactions or enzymatic processes can be attributed to these pre-steady-state signals. Moreover, the steady-state currents reflect full-cycle transport activity and correlate strongly with other measures of enzyme turnover like ATPase activity (**Fig. 3F**).

To evaluate the potential utility of these recordings to pharmacological studies, we characterized the inhibitory effect and reversibility of a known blocker of polyamine transport, AMXT-1501. This small molecule is currently in clinical trials for the treatment of aggressive forms of cancer (20-22) and was recently shown to act, at least in-part, through blockade of ATP13A2 and ATP13A3 (33, 34). Our results demonstrate AMXT-1501 inhibits polyamine-induced currents elicited from both transporters, with ATP13A3 requiring higher concentrations for complete blockade (**Fig. S8**). Since AMXT-1501 is a SPM-derived inhibitor, the lower apparent sensitivity of ATP13A3 is consistent with its preference for PUT. Furthermore, because our experimental design permits perfusion and reapplication of substrates and inhibitors, we were able to observe the notably low reversibility of AMXT-1501 in cellular membranes, with SPM-induced currents still virtually abolished after 5 minutes of washout (**Fig. 6D,E**). Indeed, the medicinal chemists who first synthesized this molecule intended the lipophilic group to anchor the molecule in the membrane, causing the SPM conjugate to act as a tethered blocker of the cellular polyamine transport machinery (32). Our SSM electrophysiological recordings demonstrate this membrane interaction, showing that AMXT-1501 but not SPM interacts with biomimetic membranes (**Fig. 6G-H**). This supports a model where persistent inhibition arises from dual transporter-membrane interactions of AMXT-1501. Future structural studies could determine how AMXT-1501 engages with the membrane in a transporter-bound structure. Importantly, these measurements serve as proof-of-concept that this electrophysiological platform can be used to evaluate the potency, transporter selectivity, and reversibility of small molecule modulators of polyamine transport in live cells.

In conclusion, we directly measured polyamine transport currents mediated by ATP13A2 and ATP13A3, demonstrating that these transporters are electrogenic and establishing a method to interrogate their function directly within live cell membranes. These recordings additionally reveal distinct components of the transport cycle and permit dynamic interrogation of how biochemical factors and small molecules modulate their activity, showing that these measurements provide information beyond what is accessible through conventional endpoint assays. Future studies may implement this platform to study polyamine transport regulatory mechanisms, evaluate polyamine transport modulating small molecules, and determine how molecular perturbations, like those caused by PARK9 variants of ATP13A2, alter specific steps of the polyamine transport cycle.

## Materials and Methods

### Plasmid constructs

The full length human ATP13A2 (isoform 1) was purchased from Epoch and subsequently subcloned into BacMam and pGEMHE vectors for expression in mammalian cells and for generating coding RNA (cRNA) for expression in oocytes, respectively. A hemagglutinin (HA) tag (YPYDVPDYA) was inserted after the valine residue at position 1030 using PCR based methods using the Phusion High Fidelity PCR kit (New England Biolabs). Similarly, the D513N mutation was incorporated using PCR primers with the QuikChange II XL Site-Directed Mutagenesis kit (Agilent). The full length human ATP13A3 (WT + D498N) were purchased from Addgene and similarly subcloned into the pGEMHE vector for generating cRNA for expression in oocytes.

### Oocyte isolation, injection, and incubation

Whole ovaries from *Xenopus laevis* (Xenopus 1, Michigan, USA) were mechanically dissected into smaller fragments, enzymatically digested with collagenase IV (Worthington, NJ, USA), and stored in ND96 solution (96 mM NaCl, 2 mM KCl, 1 mM MgCl2, 1.8 mM CaCl2, 5 mM HEPES, 50 mg/L gentamicin, pH 7.6) at 12°C. Coding RNA (cRNA) was synthesized using the mMESSAGE mMACHINE T7 Ultra kit (Thermo Fischer). We injected defolliculated stage V oocytes with 23 ng of cRNA in water and subsequently incubated the oocytes for 3 days at 12°C in ND96 solution. To evaluate the effects of DFMO, oocytes were incubated in ND96 solution supplemented with 1 mM DFMO throughout this period.

### Surface-Labeling assay in Xenopus oocytes

Three days post-injection, oocytes were blocked to prevent nonspecific binding in ND96 containing 1% bovine serum albumin (BSA) for 30 minutes at 4°C. Oocytes were then stained with 1 µg/mL rat anti-HA antibody (Roche, 3F10) in 1% BSA for 1 hour at 4°C, washed with 1% BSA, stained with HRP-conjugated anti-rat secondary antibody (Jackson ImmunoResearch, 1:1000 in 1% BSA) for 1 hour, and then washed again. Oocytes were subsequently transferred to a 96-well plate (1 oocyte/well) with wells containing ND96 without BSA. Chemiluminescent substrate (ThermoFisher, 37070) was added to the wells (100 µL/well). The plate was agitated for 1 minute and then luminescence of the wells was quantified using a Perkin Elmer Victor X3 microplate reader. Luminescence was assessed for 20 or more oocytes per condition from 3 independent experiments and differences were evaluated by 1-way ANOVA with Dunn’s *post-hoc* test.

### Two-Electrode Voltage Clamp Electrophysiology

Three days post-injection, oocytes incubating in ND96 were moved to 4°C storage until the time of recordings. For two-electrode voltage clamp (TEVC) recordings were made with an Oocyte Clamp amplifier (OC-725C; Warner Instruments), a Digidata 1550 board, a Minidigi 1B, and Clampex software (Molecular Devices, Version 10.4.0.36). Signals were filtered at 1 kHz and digitized at 20 kHz. Current and voltage microelectrodes had resistances of 0.3 to 2 MΩ when filled with 3 M KCl. During recordings, oocytes were perfused with an external bath solution containing 110 mM Na-Mes, 0.3 mM niflumic acid, 5 mM HEPES (pH 7.6), 5 mM BaCl_2_, 2 mM NiCl_2_, 2 mM MgCl_2_ hexahydrate, and 5 mM NaCl. Additional substrates and inhibitors were added as indicated. Pre-steady-state currents were determined by highlighting a range of at least 50 data points around the peak and automatically outputting the negative going peak in Clampfit software (Molecular Devices, Version 10.4.2.0). The steady-state currents were quantified by averaging a ∼30 ms range of data points at the end of the pulse. Analytical quantification of these currents was performed on raw signal traces. Hill fits of the dose-response curves was performed using Origin Software (2022b). Differences in current parameters and apparent *K*m values were analyzed using paired or unpaired two-tailed *t*-tests, one-way ANOVA, or Kruskal–Wallis tests with Dunn’s post hoc comparisons, as appropriate. The statistical test used for each analysis is specified in the corresponding figure legend. The representative signal traces shown in figures were low-pass filtered at 0.5-1 kHz.

### Western blotting

Oocyte sample lysates were prepared as previously described (44). Briefly, 5 oocytes per condition were transferred to a 1.5 mL tube, washed twice with ND96, and then homogenized by pipetting up and down 10-20 times in 100 µL of a pre-cooled solution containing 1% Triton-X, 100 mM NaCl_2_, 20 mM Tris-HCl pH 7.4, and protease inhibitors (Santa Cruz, sc-506202). Samples were subsequently centrifuged at 10,000 x g for 5 minutes at 4°C, separating the lysates into three layers: yolk, supernatant, debris (from top to bottom). A 70 µL volume of supernatant was transferred to a fresh tube, mixed with 70 µL of 4x Laemmli sample buffer (BioRad, 1610747) with β-mercaptoethanol (10%), and boiled at 95°C for 5 minutes. Samples were flash-frozen on dry ice and stored at -80°C until the assay date.

Samples were separated by molecular weight by sodium dodecyl sulfate gel electrophoresis using 4-20% Tris-Glycine pre-cast gels (Invitrogen, XP00100BOX) and then transferred to a nitrocellulose membrane. Samples were blocked for 1 hour at room temperature in a blocking solution containing 5% blotting-grade blocker (BioRad, 1706404) in TBS with Tween 20 (TBS-T) before immunostaining. Blots were stained with primary antibody in diluted in blocking solution for 2 hours at room temperature: rabbit anti-ATP13A2 (Sigma, A3361, 1:500), rat anti-HA (Roche, 3F10, 1:500), or mouse anti-tubulin (abcam, ab7291, 1:5000). Membranes were washed with TBST, stained for 1 hour with HRP secondary antibody targeted to the appropriate species (Jackson ImmunoResearch, 1:3000), and then washed again. Luminol-based substrate (ThermoFisher, 34075) was used to detect HRP stained bands by chemiluminescence using a FluorChem M System (ProteinSimple).

### Baculoviral Production and Transduction

WT- and D513N-ATP13A2-GFP baculoviruses were produced as previously described (45). Briefly, baculoviruses were generated using the Bac-to-Bac system. Plasmid constructs in the pEG BacMam expression vector were transposed into bacmids in DH10Bac cells. The resulting bacmids were transfected into Sf9 cells cultured in SF900II SFM medium. P3 baculoviruses were used to transduce HEK293S cells (1.5E6 cells/mL) cultured in 25 mL of Freestyle 293 medium for expression of ATP13A2 constructs. After 24 hours at 37°C, the infected cultures were supplemented with 10 mM sodium butyrate and grown for an additional 24 hours at 30°C before harvesting.

### Lysosomal Membrane Fraction Isolation

Lysosomal membrane fractions were isolated as previously described (46). Briefly, two hours prior to harvesting, the culture flasks containing virally transduced HEK293S were inoculated with 1 µM vacuolin to induce swelling and fusion of lysosomes into larger vesicles. Culture flasks were then returned to 30°C until harvest. Then, cells were centrifuged 500 x *g* for 10 minutes at 4°C. The supernatants were discarded, and pellets were resuspended in 1 mL pre-cooled homogenization solution (250 mM sucrose, 10 mM Tris-HCl pH 7.4 with protease inhibitor cocktail [ChemCruz, sc-29131]). Re-suspended cells were transferred to a Potter-Elvehjem homogenizer and homogenized by applying 20 strokes rapidly using a Teflon pestle. Homogenates were transferred to a 1.5 mL microcentrifuge tube and centrifuged at 14,000 x *g* for 15 minutes at 4°C. The supernatant fractions were collected, transferred to a fresh tube, and diluted with an equal volume of 16 mM CaCl_2_ to precipitate lysosomes. Diluted fractions were placed on a shaker set at 300 rpm for 5 minutes at 4°C. These fractions were then centrifuged at 25,000 x *g* for 15 minutes at 4°C. The resulting supernatants were discarded, the pellets were resuspended in wash solution (150 mM KCl, 10 mM Tris-HCl pH 7.4 with protease inhibitor cocktail [ChemCruz, sc-29131]), and then centrifuged again at 25,000 x *g* for 15 minutes at 4°C. The final pellets were resuspended in 40 µL of the wash solution and stored on ice until the activity assays or flash frozen and transferred to -80°C for long term storage.

### ATPase Activity Assays

ATPase activity of ATP13A2 expressing lysosomal fractions was determined using a commercially available assay kit (ADP-Glo Max assay, Promega) that detects the ADP produced in an ATP-consuming reaction via a luciferase luminescence-based detection assay. Protein concentrations of freshly isolated lysosomal membrane fractions were determined by bicinchoninic acid assay. ATPase assays were initiated by diluting 20 µg of lysosomal fractions in 25 µL of a reaction solution consisting of 5 mM ATP, 2 mM EGTA, 50 mM MOPS, 100 mM KCl, 11 mM MgCl_2_, 195 µM n-dodecyl-β-maltoside, 1 mM dithiothreitol, 2.1 mM CaCl_2_, and a range of SPM concentrations in a 96 well plate. Reactions were carried out over 60 minutes at 37°C. The assay was then terminated by adding 25 µL of ADP-Glo Reagent and incubating the 96 well plates on a rocker at room temperature for 40 minutes. ADP in the reactions was detected by adding 50 µL of ADP Glo MAX Detection Reagent and incubating on a rocker at room temperature for 60 minutes. The luminescent signal from the assay plate was measured using a Perkin Elmer Victor X3 microplate reader. The dose-dependence of stimulated ATPase activity was fit with a Hill equation using Origin (2022b) software.

### Solid-Supported Membrane Electrophysiology

The solid supported membrane (SSM) consisted of an octadecanethiol monolayer covalently attached to a gold electrode via the sulfur atom, with a diphytanoylphosphatidylcholine monolayer on top of the octadecanethiol layer (35). The SSM was mounted in a cuvette made from Plexiglas and acted as the working electrode. The cuvette also contained a reference electrode which was separated from the main fluid pathway by a salt bridge (47, 48). The cuvette and the fluid pathway were enclosed in a Faraday cage.

All SSM-based electrophysiological measurements were performed using a surface electrogenic event reader (SURFE^2^R) (Nanion Technologies, Munich, Germany). A detailed description of the SURFE^2^R technology can be found in the literature (47-53).

The detection of ion binding to the SSM membrane is based on the principle that the SSM behaves like an electrical capacitor, with typical values of 0.2-0.4 µFcm^-2^ for the SSM capacitance. Thus, if cations bind to the SSM membrane, electrons must flow through the external circuit towards the gold surface. Conversely, if anions bind to the SSM membrane, electrons must flow through the external circuit away from the gold electrode. In both cases a current signal is produced in the external circuit which one can measure (37, 54). When ions bind to the SSM membrane, this produces a capacitive current signal, which first rises and then decays as the ion binding reaction comes into equilibrium. The experiments were carried out by flowing an activating solution containing either SPM or AMXT-1501 in buffer (100 mM NaCl, 10 mM HEPES, pH 7.6) across the SSM membrane, followed by a wash with non-activating solution, i.e., buffer without SPM or AMXT-1501. The temperature was maintained at 20-22 °C for all the experiments. To verify the reproducibility of the current signals on the same SSM membrane, each single measurement was repeated five times (1 minute time interval between measurements) and then averaged to improve the signal-to-noise ratio. Standard deviations did not exceed 5%. Moreover, each set of measurements was usually reproduced using two different SSM membranes.

## Acknowledgements

The authors thank Deepa Srikumar for technical assistance and logistical support. We would also like to thank Dr. Joseph A. Mindell’s laboratory for generously sharing the LAMP1 antibody and access to their ultracentrifugation and plate reader instruments. The authors would also like to thank Drs. Natalie Landeck and Antoniya Aleksandrova for helpful discussions. The cartoon depictions of ATP13A2 were created using BioRender. This research was supported in part by the Intramural Research Program of the National Institutes of Health (NIH). The contributions of the NIH authors were made as part of their official duties as NIH federal employees, are in compliance with agency policy requirements, and are considered Works of the United States Government. However, the findings and conclusions presented in this paper are those of the author(s) and do not necessarily reflect the views of the NIH or the U.S. Department of Health and Human Services. M.H. is supported by the Intramural Research Program of the National Institute of Neurological Disorders and Stroke (NINDS), NIH (ZIANS002993). S.R.C. is supported by an NINDS Competitive Postdoctoral Fellowship Award through the Intramural Research Program.

## Notes

### Competing Interest Statement

The authors have declared no competing interest.

