## Supplementary material for "Direct detection of electrogenic polyamine transport": Fig. S1

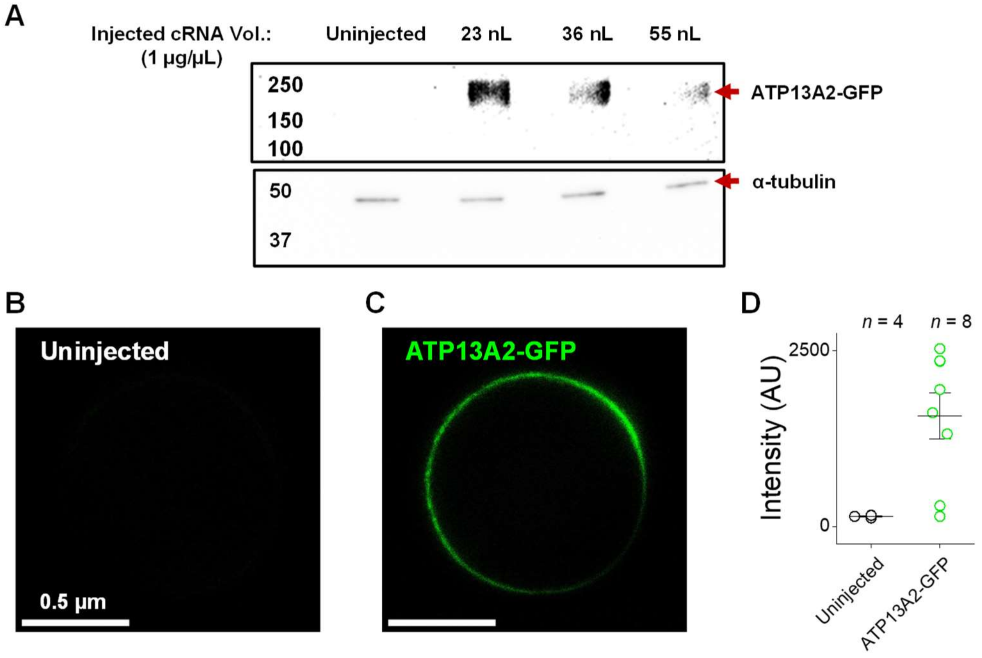


**Supplementary Figure S1. A)** Western blot of lysates from oocytes injected with ATP13A2-GFP cRNA immunostained with anti-ATP13A2 and -alpha-tubulin antibodies, shows that injecting lower volumes of cRNA (1 µg/µL) resulted in higher ATP13A2 expression. **B, C)** Confocal image of uninjected (*B*) or ATP13A2-GFP injected (*C*) oocytes taken with a 10X objective. **D)** Quantification of GFP fluorescence intensity of uninjected oocytes and oocytes injected with ATP13A2-GFP cRNA.
