## Supplementary material for "Direct detection of electrogenic polyamine transport": Fig. S2

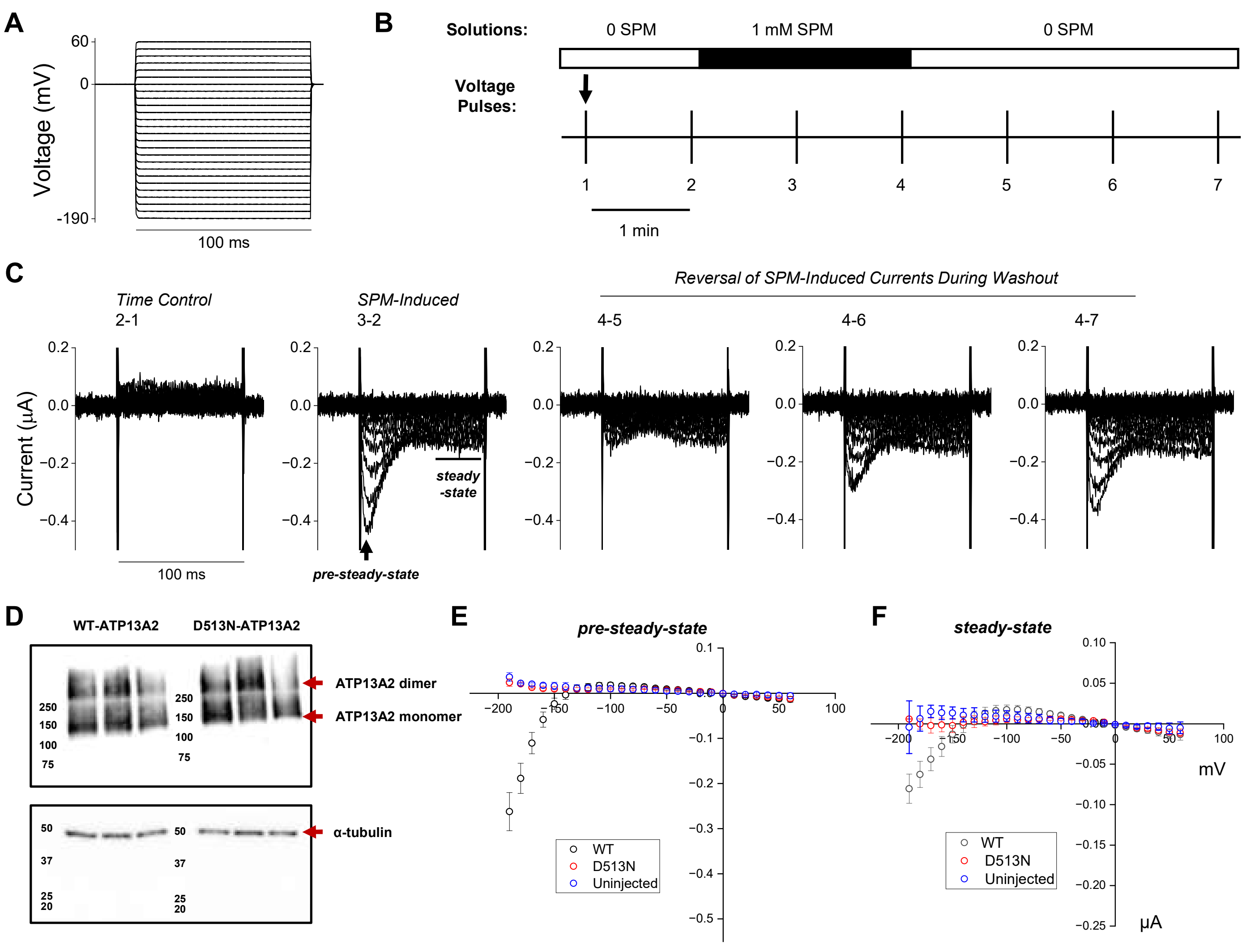


**Supplementary Figure S2. A,B)** A voltage protocol of 100 ms pulses from -190 to +40 mV (10 mV step, holding potential = 0 mV) was applied to oocytes every 1 min while perfusing with bath solution ± 1 mM SPM. **C)** SPM-induced currents were detected within 1 minute of substrate addition (P3-P2), and the steady-state currents were reversed within 3 minutes of perfusion with bath solution (P4-P7). **D)** Lysates of oocytes injected with WT-(*left*) or D513N-(*right*)ATP13A2-GFP cRNA were taken after 3 independent TEVC experiments to confirm expression of the constructs after recordings. Immunostaining with anti-ATP13A2 and -alpha tubulin antibodies shows both WT and catalytically incompetent constructs are well expressed in oocytes. **E, F)** Voltage-dependence of pre-steady-state (*E*) and steady-state (*F*) currents elicited from oocytes injected with ATP13A2-WT (*black*, *n* = 14) ATP13A2-D513N (*red*, *n* = 7), or uninjected oocytes (*blue*, *n* = 3).
