## Supplementary material for "Direct detection of electrogenic polyamine transport": Fig. S3

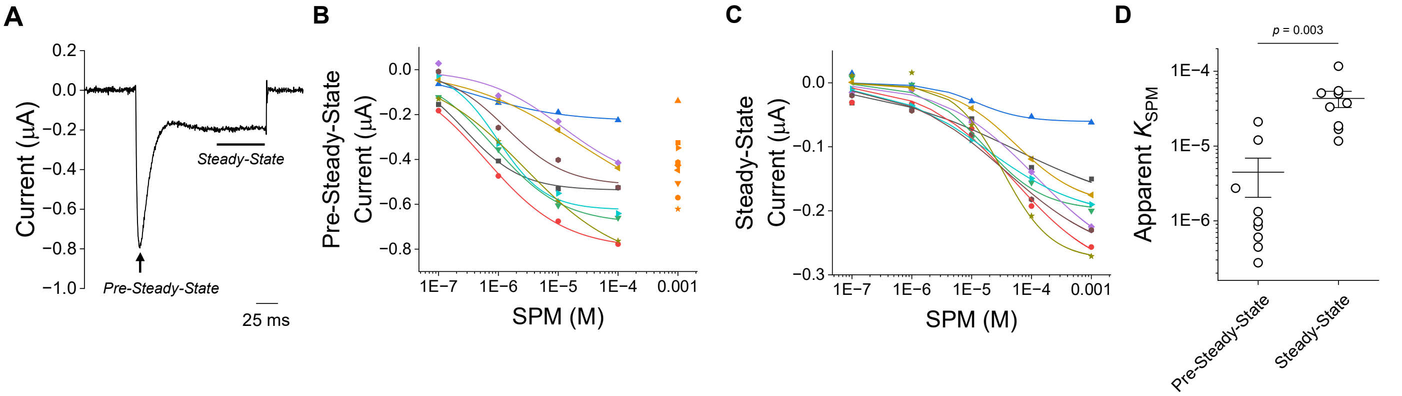


**Supplementary Figure S3. A)** Representative SPM-induced currents elicited from ATP13A2 injected oocytes during a 150 ms voltage step from 0 to -190 mV. **B, C)** Dose-dependent activation pre-steady-state (*B*) and steady-state currents (*C*) ($n = 9$). Data from each independent oocyte were fit with a Hill equation. For the pre-steady-state component, only the ascending phase was fit with Hill equation. The orange points at high SPM concentration were therefore excluded from the fits. **D)** Apparent $K_{SPM}$ was significantly lower for the pre-steady-state component compared to the steady-state ($p = 0.004$, two-tailed paired t-test).
