## Supplementary material for "Direct detection of electrogenic polyamine transport": Fig. S4

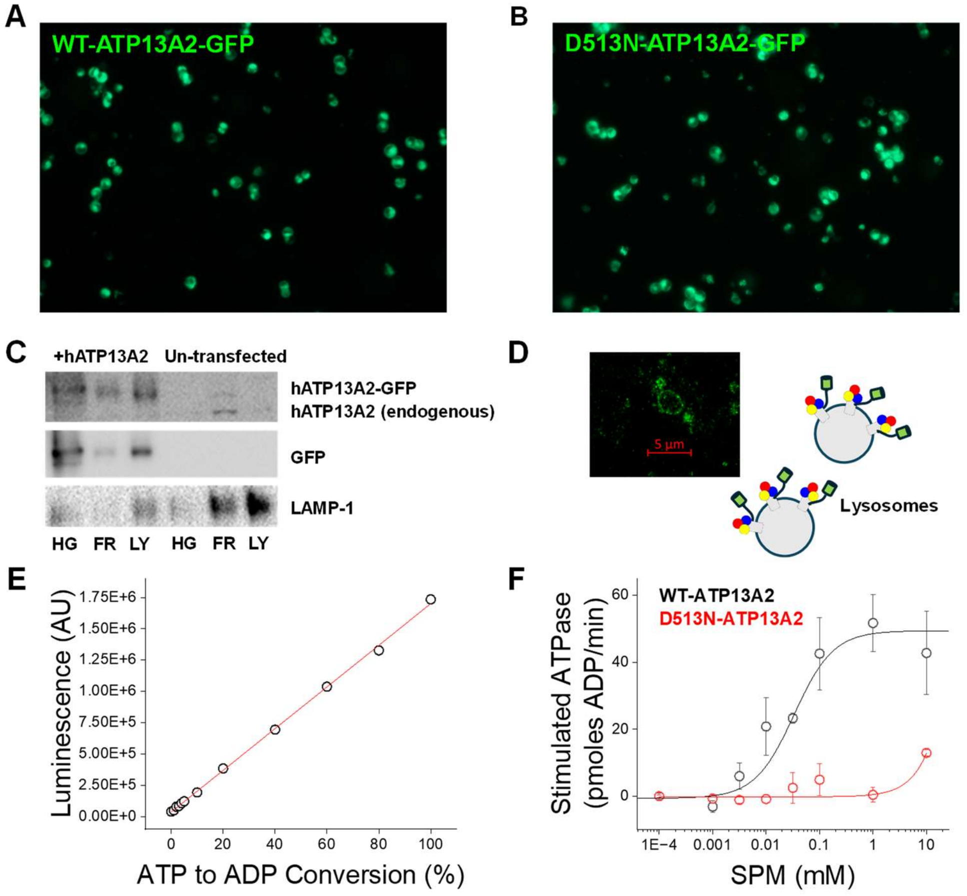


**Supplementary Figure S4. A,B)** HEK293S cells expressing WT- (A) or D513N- (B) ATP13A2-GFP after baculoviral transduction. **C)** Western blot demonstrates that ATP13A2-GFP and LAMP-1 are co-enriched as the samples are purified from the starting cellular homogenates (HG) to organellar membrane fractions (FR), and to lysosomal fractions (LY). **D)** Confocal image of lysosomes after they were enlarged by pre-treating HEK293 cells with 1 µM vacuolin and isolated through homogenization and centrifugation. **E)** A standard curve was used to estimate ATP to ADP conversion based on ATP-dependent activation of luciferase luminescence. **F)** Stimulated ATPase activity of lysosomal fractions from cells expressing WT-ATP13A2-GFP (black) or D513N-ATP13A2-GFP (red). Data are mean ± SEM ($n$ = 3). Stimulated ATPase activity from WT-ATP13A2 was well described by a Hill equation (apparent $K_m$ = 33.4 ± 12.7 µM, *Hill coefficient* = 1.2 ± 0.6, $R^2$ = 0.971).
