## Supplementary material for "Direct detection of electrogenic polyamine transport": Fig. S5

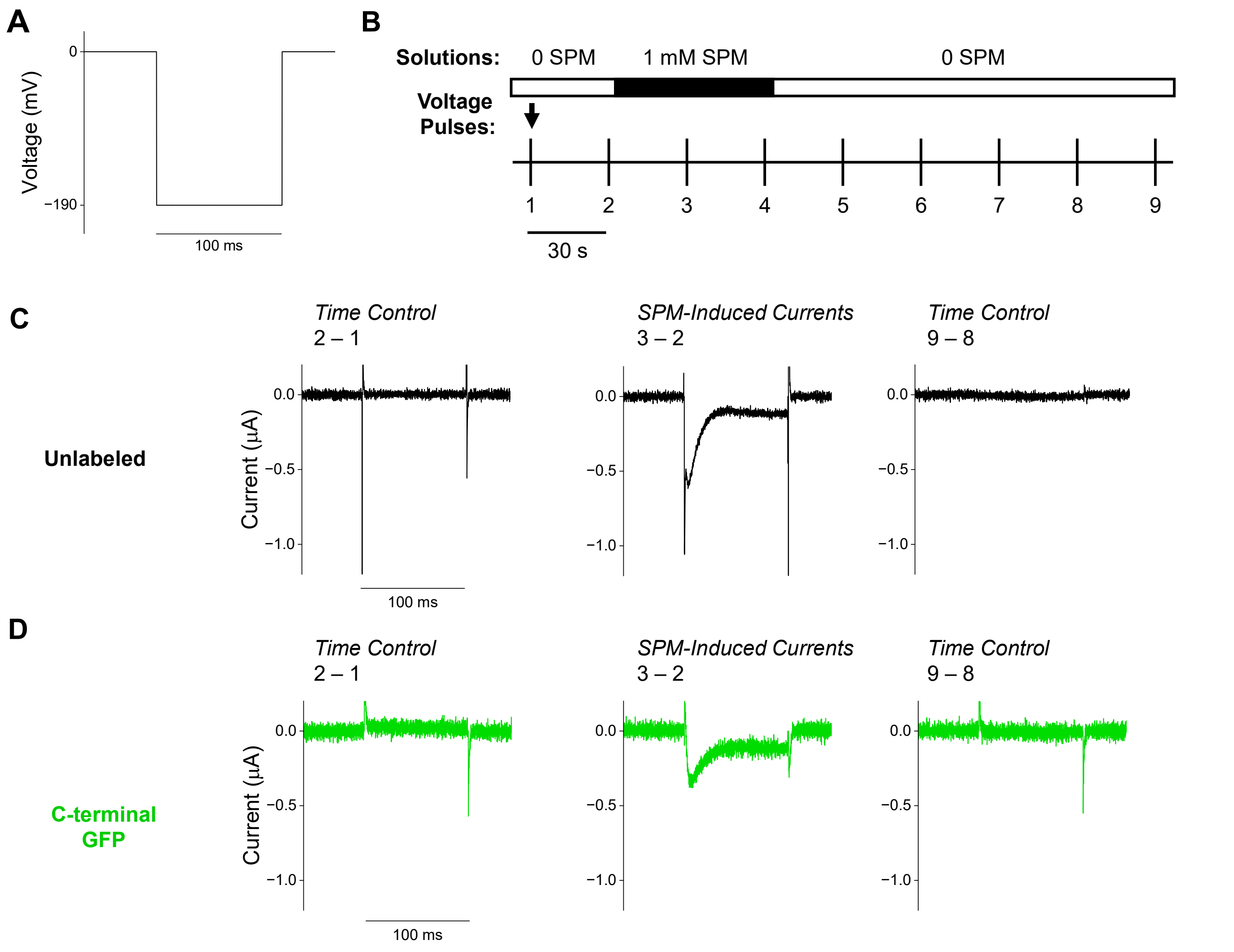


**Supplementary Figure S5. A, B)** Voltage steps from 0 to -190 mV (100 ms pulse, holding potential = 0 mV) were applied to oocytes every 30 seconds while perfusing with bath solution ± 1 mM SPM. **C,D)** Similar currents were elicited from oocytes injected with unlabeled- (*C*) or C-terminal GFP-labeled- ATP13A2 cRNA (*D*) within the same batch.
