## Supplementary material for "Direct detection of electrogenic polyamine transport": Fig. S6

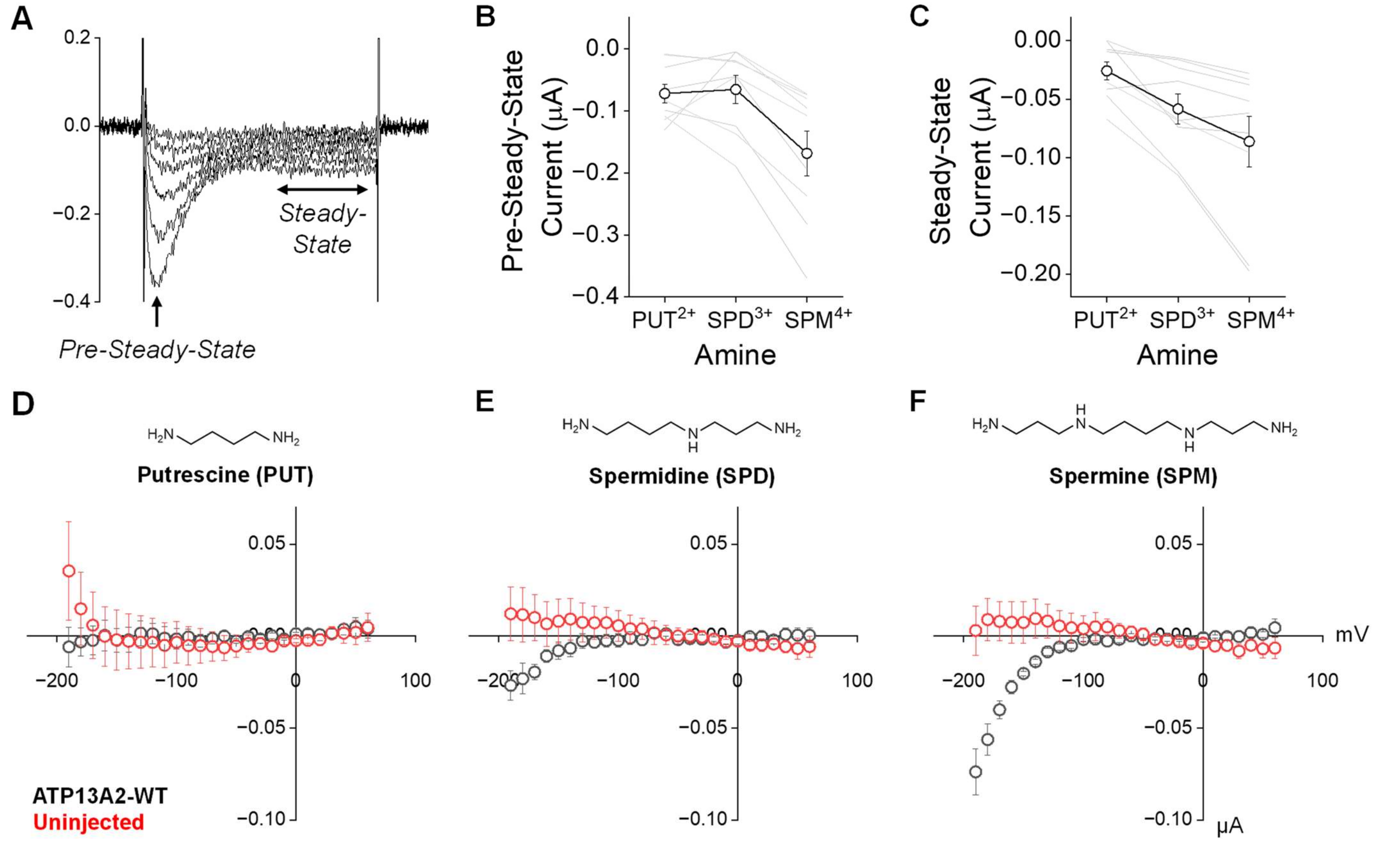


**Supplementary Figure S6. A)** Representative traces of currents elicited from ATP13A2 injected oocytes at -140 to -190 mV. **B, C)** Polyamine substrate-dependent activation pre-steady-state (*B*) and steady-state currents (*C*) ($n = 9$). **D-F)** Voltage-dependences of polyamine-induced currents activated by 1 mM PUT (*D*), SPD (*E*), and SPM (*F*) in oocytes injected with ATP13A2-WT cRNA (*black*, $n = 4$) or uninjected oocytes (*red*, $n = 3$).
