## Supplementary material for "Direct detection of electrogenic polyamine transport": Fig. S7

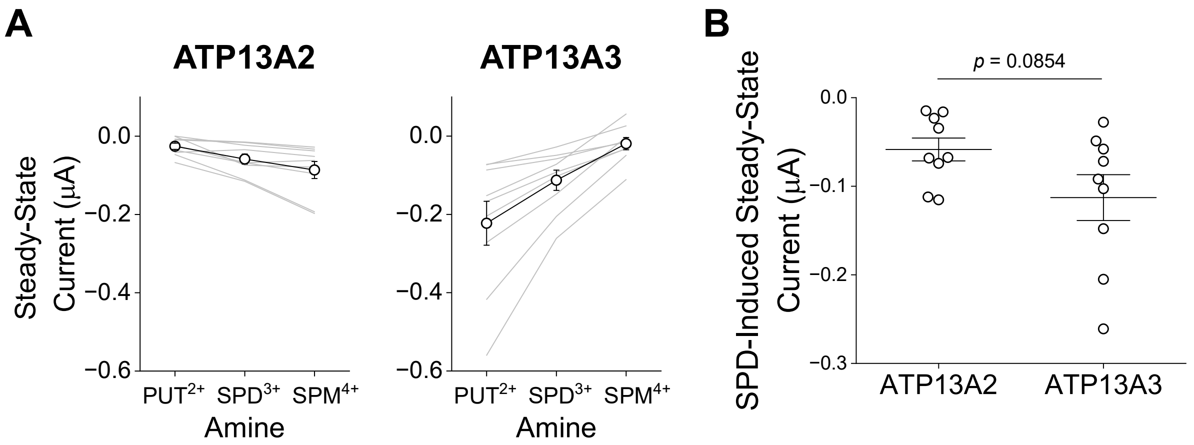


**Supplementary Figure S7. A)** Steady-state currents elicited from ATP13A2 and ATP13A3 induced by 1 mM PUT, SPD, or SPM in response to voltage pulses from 0 to -190 mV (100 ms pulse, holding potential = 0 mV) reflect the known substrate preferences of these transporters. **B)** SPD-induced (1 mM) steady-state currents at -190 mV appeared to be larger for ATP13A3 than ATP13A2, although the difference was not statistically significant ($p = 0.0854$, two-tailed t-test).
