## Supplementary material for "Direct detection of electrogenic polyamine transport": Fig. S8

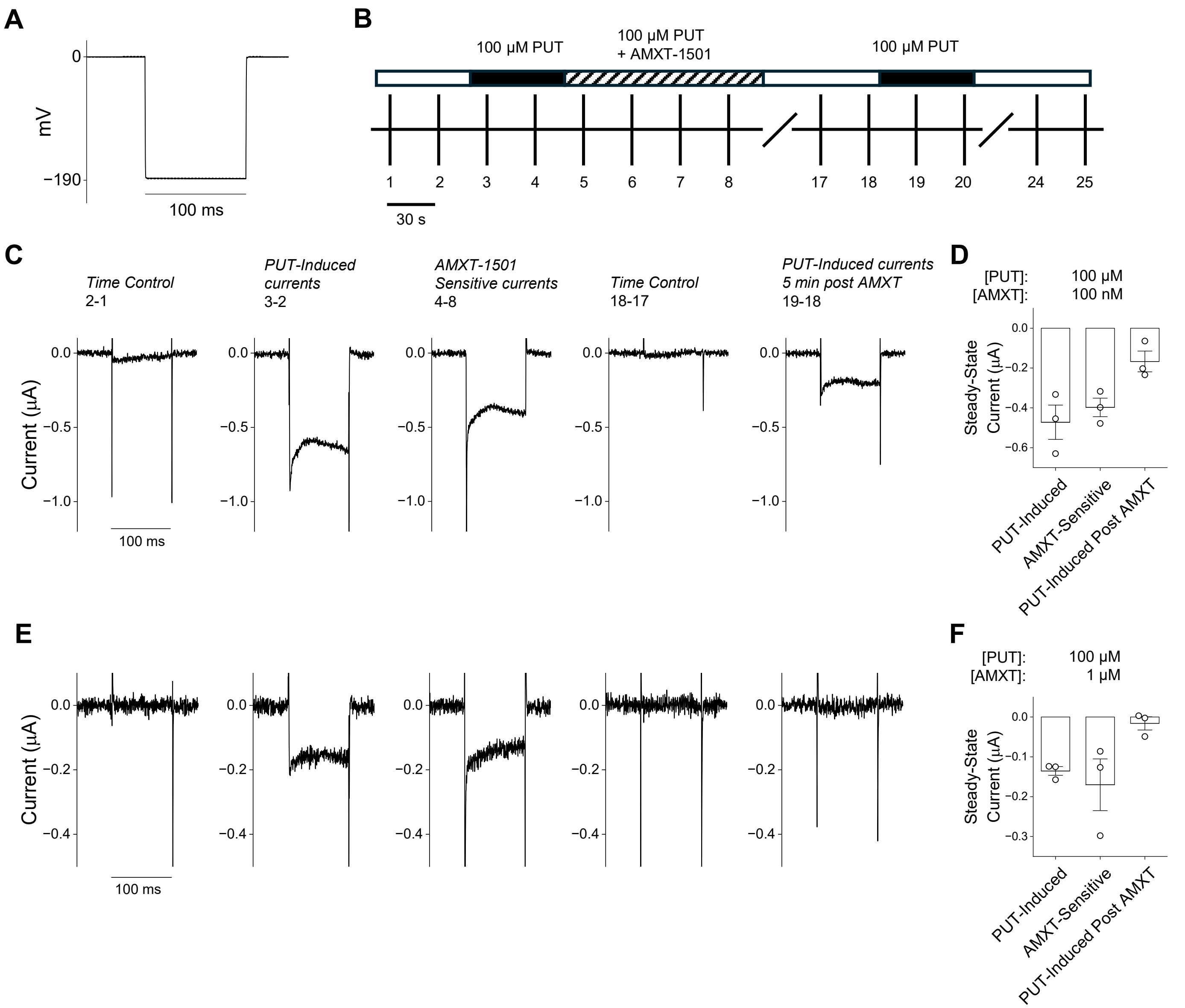


**Supplementary Figure S8. A, B)** -190 mV, 100 ms depolarizations (holding potential = 0 mV) were applied to ATP13A3-injected oocytes every 30 s while perfusing with bath solution or solution with 100 µM PUT ± a concentration of AMXT-1501. **C,D)** Only a fraction of the current induced by 100 µM PUT was blocked by 100 nM AMXT-1501. Some current was recovered after removing AMXT-1501 and perfusing with bath solution for 5 min. (*n* = 3). **E,F)** 1 µM AMXT-1501 completely abolished PUT-induced currents. PUT-induced currents were not recovered after removing AMXT-1501 and perfusing with bath solution for 5 min (*n* = 3).
