## Supplementary material for "Direct detection of electrogenic polyamine transport": Fig. S9

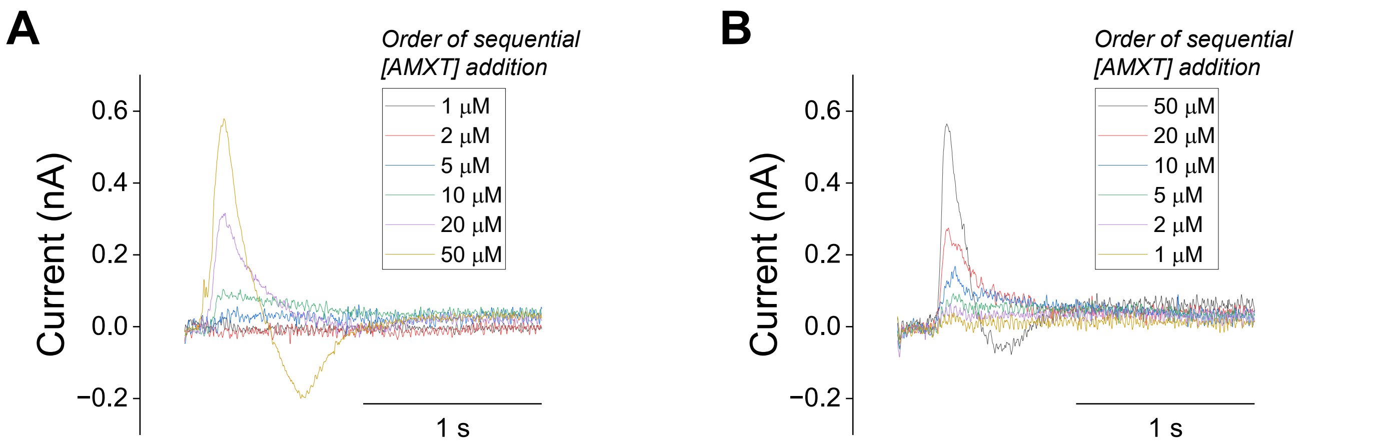


**Supplementary Figure S9. A,B)** Solid-supported membrane current traces were recorded following AMXT-1501 concentration jumps. The same concentration jumps were performed in ascending (*A*) and descending (*B*) AMXT-1501 concentration order, as indicated in the panels. Each trace was obtained by averaging five consecutive measurements ($n$ = 5).
